# Durable Immunity to Ricin Toxin Elicited by a Thermostable, Lyophilized Subunit Vaccine

**DOI:** 10.1101/2021.09.08.459551

**Authors:** Hayley Novak, Jennifer Doering, Dylan Ehrbar, Oreola Donini, Nicholas J. Mantis

## Abstract

The development of vaccines against biothreat toxins like ricin (RT) is considered an integral component of the United States national security efforts. RiVax^®^ is a thermostable, lyophilized RT subunit vaccine adsorbed to aluminum salt adjuvant intended for use by military personnel and first responders. Phase 1 studies indicated that RiVax is safe and immunogenic, while a three dose, intramuscular vaccination regimen in non-human primates elicited protection against lethal dose RT challenge by aerosol. Here we investigated, in a mouse model, the durability of RiVax-induced antibody responses and corresponding immunity to lethal dose RT challenge. Groups of mice were subcutaneously administered 3 or 1 μg of RiVax on days 0 and 21 and challenged with 10 × LD_50_ RT by injection at six different intervals over the course of twelve months. Serum antibody titers and epitope-specific competition assays were determined prior to each challenge. We report that the two-dose, 3 μg regimen conferred near complete protection against RT challenge on day 35 and complete protection thereafter (challenge days 65, 95, 125, 245, and 365). The two-dose, 3 μg regimen was superior to the 1 μg regimen as revealed by slight differences in survival and morbidity scores (e.g., hypoglycemia, weight loss) on challenge days 35 and 365. In separate experiments, a single 3 μg RiVax vaccination proved only marginally effective at eliciting protective immunity to RT, underscoring the necessity of a prime-boost regimen to achieve full and long-lasting protection against RT.

**IMPORTANCE:** Ricin toxin (RT) is a notorious biothreat, as exposure to even trace amounts via injection or inhalation can induce organ failure and death within a matter of hours. In this study, we advance the preclinical testing of a candidate RT vaccine known as RiVax^®^. RiVax is a recombinant non-toxic derivative of RT’s enzymatic subunit that has been evaluated for safety in Phase I clinical trials and efficacy in a variety of animal models. We demonstrate that two doses of RiVax is sufficient to protect mice from lethal dose RT challenge for up to one year. We describe kinetics and other immune parameters of the antibody response to RiVax and discuss how these immune factors may translate to humans.

## Introduction

Ricin toxin (RT) is classified by the Centers for Disease Control and Prevention (CDC) as a biothreat toxin, alongside botulinum neurotoxin (BoNT), abrin, and *Staphylococcus* enterotoxin (SEB) (1). A recent NATO commission ranked RT as having a particularly high threat potential owing to its toxicity via inhalation, its ease of acquisition, and a lack of available countermeasures (2). RT is a product of the castor bean, *Ricinus communis*, which is propagated globally for its oils that are used in lubricants and cosmetics (3). In its mature form, RT is a ∼65 kDa cytotoxic glycoprotein consisting of two similarly sized subunits, RTA and RTB. RTB is a galactose/N-acetyl galactosamine (Gal/GalNAc)-specific lectin that facilitates RT entry into all mammalian cell types, including alveolar macrophages and airway epithelial cells (4). Once inside a cell, RTA is a highly efficient ribosome-inactivating protein (RIP) that triggers cell death and inflammation within a matter of hours (5-8). When inhaled, RT evokes a form of respiratory distress characterized by neutrophilic infiltration, intra-alveolar edema, accumulation of pro-inflammatory cytokines like IL-1 and IL-6 in bronchoalveolar lavage (BAL) fluids, and fibrinous exudate (9, 10). Monoclonal antibody (MAb) intervention studies in non-human primates indicate that the “point of no return” following a RT aerosol exposure is on the order of hours (11).

One of the most advanced candidate RT vaccines is RiVax^®^, a recombinant, non-toxic derivative of RTA (12-14). RiVax and RiVax-adsorbed to Alhydrogel^®^ is safe and immunogenic in healthy adults (15, 16). We have reported that RiVax is stable for at least 12 months at 40°C without loss of potency when formulated as a dry, glassy solid containing colloidal aluminum hydroxide adjuvant with trehalose as a stabilizing excipient (17, 18). In non-human primates, the thermostable RiVax formulation (100 μg) administered intramuscularly three times at monthly intervals stimulated immunity to subsequent lethal dose RT aerosol challenge (19). Protection correlates with the onset of RT-specific endpoint titers, as well as epitope-specific serum antibody responses revealed through a competition ELISA known as EPICC (D. Ehrbar, C. Roy, G. Van Slyke, E. Vitetta, and N. Mantis, manuscript in preparation) (19, 20).

In this report we sought to further advance the pre-clinical analysis of the thermostable formulation of RiVax by examining the durability of protective immunity and RT-specific antibody responses induced following vaccination in mice. We were particularly interested in a prime-boost regimen in light of recent dose-response studies performed in Rhesus macaques (D. Ehrbar, C. Roy, G. Van Slyke, O. Donini, E. Vitetta, and N. Mantis, manuscript in preparation).

## RESULTS

To assess the durability of RiVax-mediated immunity, 12 groups of mice were vaccinated SC with RiVax (3 μg; n=12 per group) or vehicle (PBS) only (n=6 per group) on days 0 and 21. Groups of mice were challenged with 10 × LD_50_ RT by IP injection on days 35, 65, 95, 125, 245, and 365 (**Table 1**). Two additional groups of mice were vaccinated SC with a low dose (1 μg) of RiVax on days 0 and 21 and challenged with RT on days 35 and 365 (**Table 1**). Serum samples were collected from animals five days prior to RT challenge and were assessed for RT-specific serum IgG endpoint titers by ELISA. The serum samples were also evaluated using an epitope-specific competition ELISA known as EPICC (20). As indicators of morbidity, we monitored animal weight loss and onset of hypoglycemia (21-23). A reference lot of RiVax was used for all experiments (18). The vaccine was reconstituted with sterile water for injection (WFI) immediately prior to use. It should be noted that, at all timepoints, groups of mice that were sham vaccinated succumbed to RT intoxication within 3 days (**Table 1**).

**Table 1.**
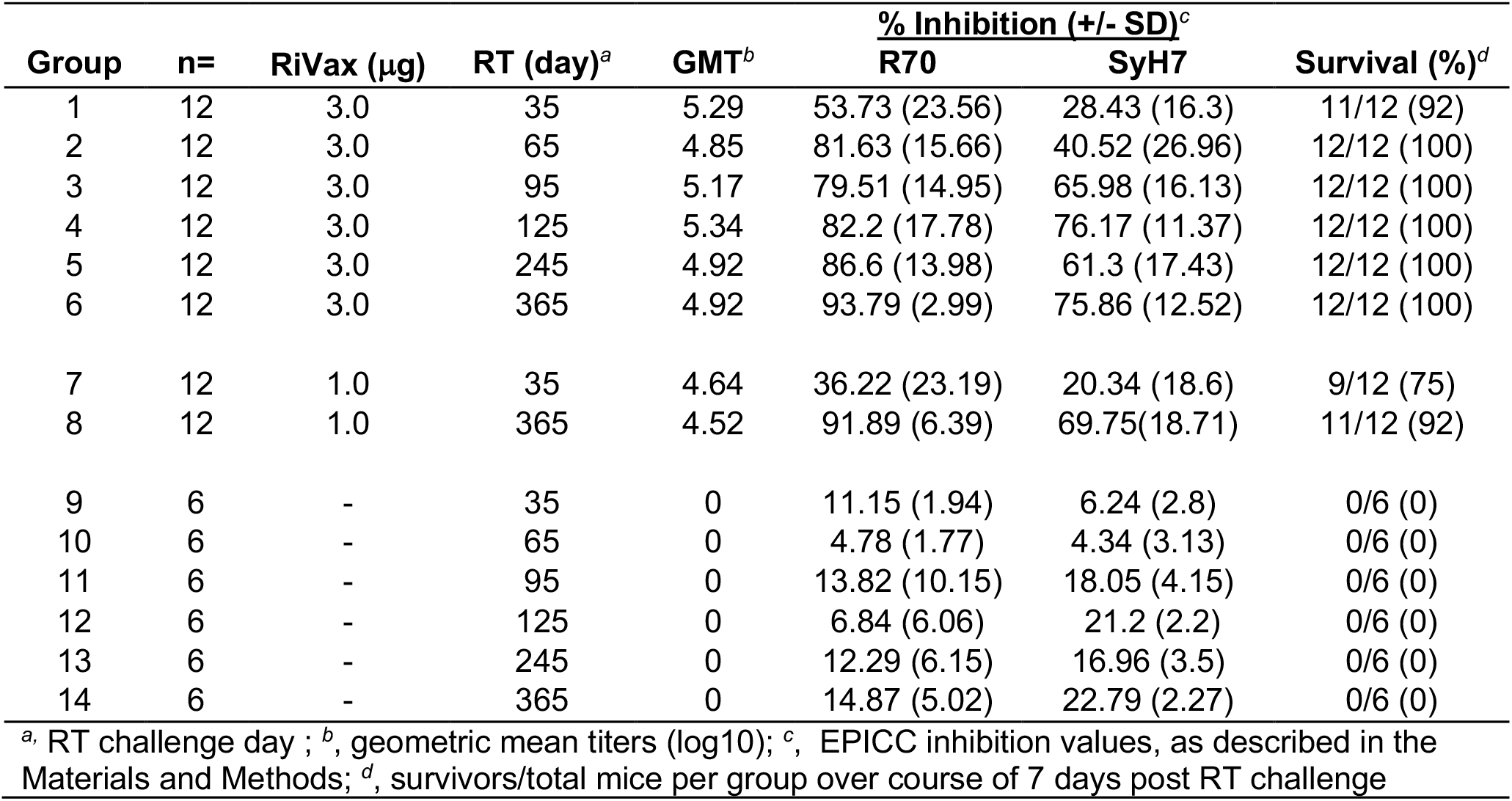
Response elicited by RiVax prime-boost vaccination regimen followed by RT challenge (IP)

| Group | n= | RiVax (μg) | RT (day)<sup>a</sup> | GMT<sup>b</sup> | % Inhibition (+/- SD)<sup>c</sup> |  | Survival (%)<sup>d</sup> |
| --- | --- | --- | --- | --- | --- | --- | --- |
|  |  |  |  |  | R70 | SyH7 |  |
| 1 | 12 | 3.0 | 35 | 5.29 | 53.73 (23.56) | 28.43 (16.3) | 11/12 (92) |
| 2 | 12 | 3.0 | 65 | 4.85 | 81.63 (15.66) | 40.52 (26.96) | 12/12 (100) |
| 3 | 12 | 3.0 | 95 | 5.17 | 79.51 (14.95) | 65.98 (16.13) | 12/12 (100) |
| 4 | 12 | 3.0 | 125 | 5.34 | 82.2 (17.78) | 76.17 (11.37) | 12/12 (100) |
| 5 | 12 | 3.0 | 245 | 4.92 | 86.6 (13.98) | 61.3 (17.43) | 12/12 (100) |
| 6 | 12 | 3.0 | 365 | 4.92 | 93.79 (2.99) | 75.86 (12.52) | 12/12 (100) |
| 7 | 12 | 1.0 | 35 | 4.64 | 36.22 (23.19) | 20.34 (18.6) | 9/12 (75) |
| 8 | 12 | 1.0 | 365 | 4.52 | 91.89 (6.39) | 69.75(18.71) | 11/12 (92) |
| 9 | 6 | - | 35 | 0 | 11.15 (1.94) | 6.24 (2.8) | 0/6 (0) |
| 10 | 6 | - | 65 | 0 | 4.78 (1.77) | 4.34 (3.13) | 0/6 (0) |
| 11 | 6 | - | 95 | 0 | 13.82 (10.15) | 18.05 (4.15) | 0/6 (0) |
| 12 | 6 | - | 125 | 0 | 6.84 (6.06) | 21.2 (2.2) | 0/6 (0) |
| 13 | 6 | - | 245 | 0 | 12.29 (6.15) | 16.96 (3.5) | 0/6 (0) |
| 14 | 6 | - | 365 | 0 | 14.87 (5.02) | 22.79 (2.27) | 0/6 (0) |

<sup>a</sup>, RT challenge day ; <sup>b</sup>, geometric mean titers (log10); <sup>c</sup>, EPICC inhibition values, as described in the Materials and Methods; <sup>d</sup>, survivors/total mice per group over course of 7 days post RT challenge

The group of mice that received 3 μg of RiVax and challenged with RT on day 35 had a 92% (11/12) survival rate (**Table 1; Figure 1**). At timepoints thereafter, the groups of mice that received high dose (3 μg) RiVax exhibited complete survival (**Table 1**; **Figure 1**). From the standpoint of morbidity, mice that were vaccinated with 3 μg of RiVax and challenged on day 35 experienced significant weight loss and hypoglycemia in the days immediately following toxin exposure, whereas mice challenged on day 365 (or any other day besides day 35) did not (**Figure 1**). Mice that received the low dose of RiVax (1 μg) exhibited 75% (9/12) survival upon RT challenge on day 35 and 92% (11/12) survival on day 365. The low-dose vaccinated mice that survived RT challenge did not experience more severe weight loss or hypoglycemia than their counterparts that received high dose RiVax (**Figure 1**). Collectively, this data demonstrates that a prime-boost regimen with RiVax (3 μg) elicits a protective immune response to RT that matures between days 35 and 60 and persists for at least 12 months.

**Figure 1:**
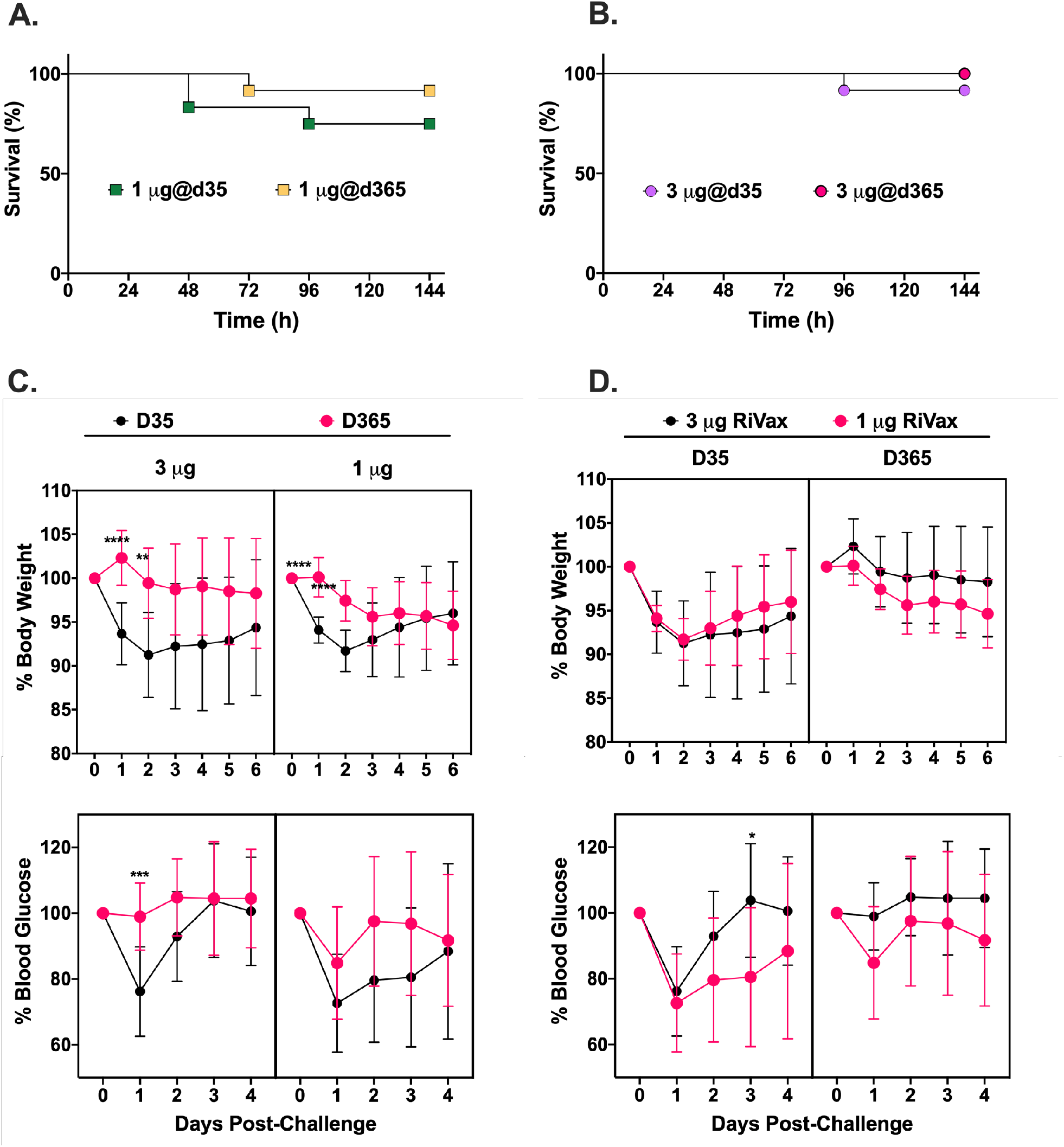
Durability of RiVax-induced immunity to ricin toxin (RT) over a period of one year. Groups of BALB/c mice were vaccinated with 1 μg or 3 μg RiVax on days 0 and 21 and then challenged with 10 × LD50 at timepoints over the course of one year, as shown in **Table 1**. Kaplan-Meier curves of groups of mice vaccinated with (**A**) 1 μg or (**B**) 3 μg RiVax and challenged with RT on day 35 or 365. Survival was monitored over a period of 7 days. (**C-D**) Body weight and blood glucose levels following RT challenge, plotted as a function of RiVax dose and day of challenge. Shown are mean values +/- SD. Significance was determined using a two-way ANOVAs followed by Šidák’s multiple comparisons tests. * P ≤ 0.05, ** P < 0.01, *** P < 0.001, **** P < 0.0001.

In light of these results, we evaluated the kinetics of RT-specific serum IgG endpoint titers in the low- and high-dose RiVax vaccinated mice (**Figure 2**). On day 30, there was a ∼5-fold difference in GMT between mice that received the 1 μg (4.442 GMT) and 3 μg (5.118 GMT) (**Table 1**; **Figure 2**). However, on day 60 and beyond, the GMTs were similar (**Table 1**; **Figure 2**), indicating the high dose of RiVax accelerated the onset of RT-specific antibody titers by day 30, but thereafter antibody levels were equivalent and remained steady for the duration of the 12-month study.

**Figure 2:**
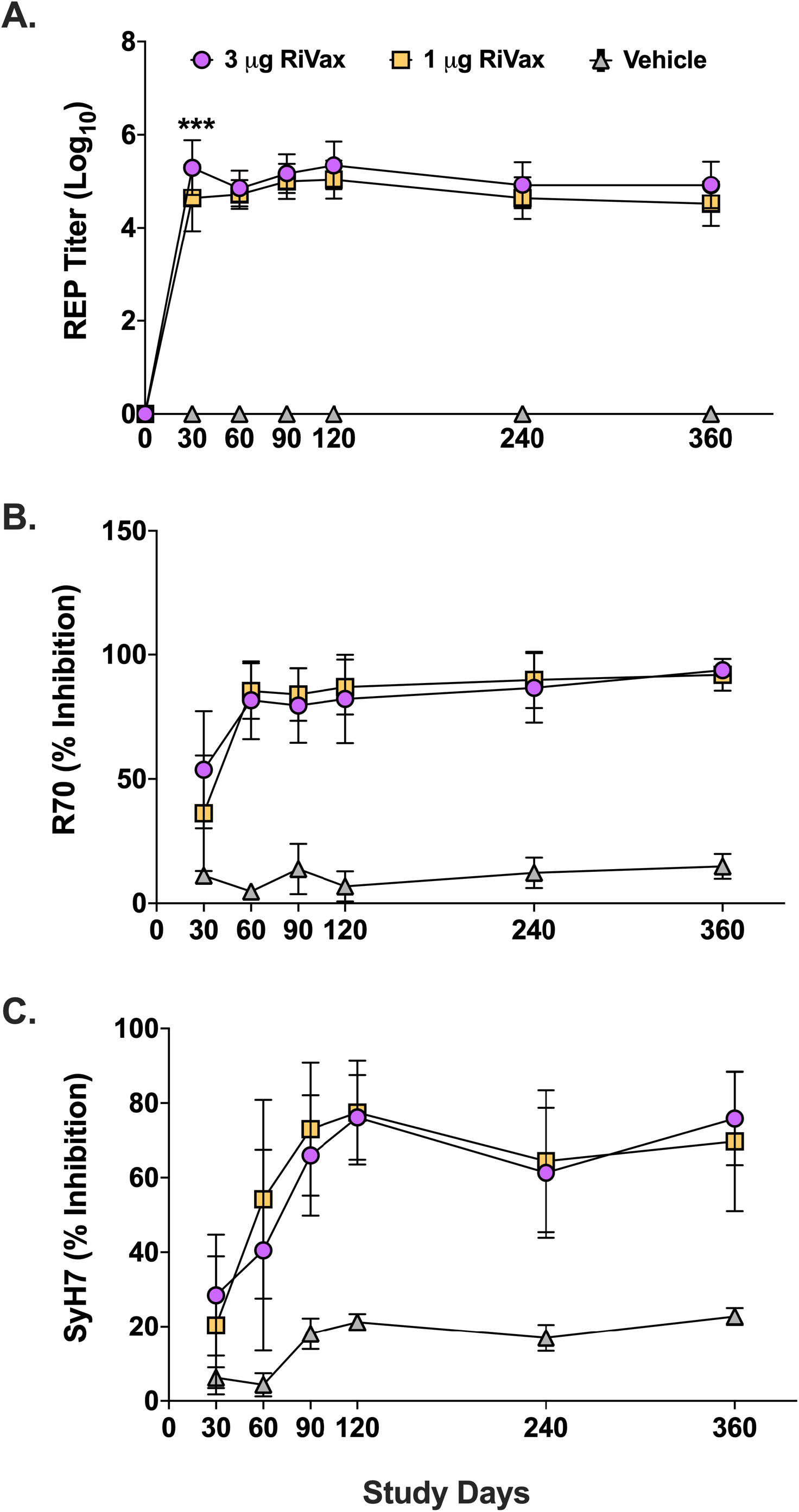
Durability of RT- and epitope-specific antibody in responses of mice following RiVax vaccination. Graphical representations of (**A**) RT-specific reciprocal endpoint (REP) titers and (**B-C**) EPICC inhibition values from the sera of mice vaccinated with 1 μg or 3 μg RiVax on days 0 and 21, as reported in **Table 1**. Sera were collected on the experimental days indicated on the x-axis. Endpoint titers were determined by ELISA using RT as coating antigen. EPICC analyses were done with (B) R70 and (C) SyH7. In all cases, mean values +/- SD are shown. Mixed effects analyses followed by Šidák’s multiple comparisons tests were used to determine significant differences between groups. * P ≤ 0.05, ** P < 0.01, *** P < 0.001, **** P < 0.0001.

We next used EPICC to assess differences in epitope-specific antibody responses over time and between the low and high RiVax doses. EPICC is a modified capture ELISA in which microtiter plates are coated with MAbs, then probed with soluble biotinylated RT, in the absence or presence of a competitor antiserum (20). R70 and SyH7 are toxin-neutralizing MAbs that recognize two spatially distinct immunodominant epitopes on RTA (24). A reduction in captured biotinylated RT by R70 and/or SyH7 is inversely proportional to the amount of competitor antibodies present in the query antisera (20). Furthermore, we have demonstrated that EPICC inhibition values serve as a correlate of vaccine-induced immunity to RT in mice (20).

We found that R70 EPICC inhibition values were detectable by day 30 in the sera of mice that received low and high dose RiVax, plateaued at day 60, and persisted for the remainder of the study (**Figure 2B**). In the 3 μg RiVax group, R70 EPICC inhibition values were slightly higher on day 30 compared to 1 μg RiVax group but were essentially identical thereafter (**Figure 2B**). SyH7 EPICC inhibition levels for both vaccination groups were low on day 30 but rose until they plateaued on day 120, reflecting a delay in maturation of the SyH7 response (**Figure 2C**). The EPICC inhibition values persisted for at least 360 days and corresponded with overall immunity to RT challenge.

### Immunity to lethal dose RT challenge elicited by a single RiVax vaccination

It has become apparent in the COVID-19 pandemic that in public health emergencies it may be necessary to implement single dose vaccinations when supplies of vaccines are limited. To investigate the benefit afforded by a single dose of RiVax (3 μg), a cohort of mice were vaccinated SC with 3 μg of RiVax on day 0. Groups of mice (n=12) were then challenged with 10 × LD_50_ RT on days 35, 65 and 95. As noted above, we monitored mice for weight loss and hypoglycemia as indicators of toxin-induced morbidity. In addition, serum samples prior to toxin challenge were evaluated by RT-specific ELISA and EPICC.

The single vaccination regimen proved vastly inferior to the two-dose regimen in terms of eliciting protective immunity to RT. Only 17% (2/12) of the mice that received the single vaccine survived RT challenge on days 35 and 65 (**Table 2**). On day 95, survival was just 33% (4/12), compared to 100% in the two-dose regimen (**Table 2**; **Figure 3**). Of note, RT-specific serum IgG levels in the mice that received the single dose of RiVax were similar on days 30, 60, and 90, revealing the absence of any maturation of antibody titers beyond day 30 (**Figure 3**). In fact, the single dose of RiVax elicited a significantly lower RT-specific serum IgG titers across the board, according to a mixed effects analysis with Šídák’s multiple comparisons test. Those same groups of mice had low EPICC inhibition values compared to the mice that received two doses of RiVax (add to **Figure 3**). These data demonstrate that, at the dose tested, little benefit is afforded by a single vaccination with RiVax.

**Table 2.** Response elicited by single RiVax vaccination regimen.

| **Table 2. Response elicited by single RiVax vaccination regimen** |  |  |  |  |  |  |  |
| --- | --- | --- | --- | --- | --- | --- | --- |
| **Group** | **n=** | **RiVax (µg)** | **RT (day)<sup>a</sup>** | **GMT<sup>b</sup>** | **% Inhibition (+/- SD)<sup>c</sup>** |  | **Survival (%)<sup>d</sup>** |
|  |  |  |  |  | **R70** | **SyH7** |  |
| 1 | 12 | 3.0 | 35 | 3.289 | 9.80 (6.07) | 12.92 (2.75) | 2/12 (17) |
| 2 | 12 | 3.0 | 65 | 3.289 | 25.24 (17.79) | 10.87 (6.91) | 2/12 (17) |
| 3 | 12 | 3.0 | 95 | 3.289 | 1.45 (19.49) | 21.82 (31.91) | 4/12 (33) |
| 4 | 6 | - | 35 | 0 | 5.98 (2.06) | 14.6 (2.83) | 0/6 (0) |
| 5 | 6 | - | 65 | 0 | 13.23 (3.46) | 8.74 (5.85) | 0/6 (0) |
| 6 | 6 | - | 95 | 0 | -12.58 (3.83) | 9.67 (26.41) | 0/6 (0) |

<sup>a</sup>, RT challenge day ; <sup>b</sup>, geometric mean titers (log10); <sup>c</sup>, EPICC inhibition values, as described in the Materials and Methods; <sup>d</sup>, survivors/total mice per group over course of 7 days post RT challenge

**Figure 3:**
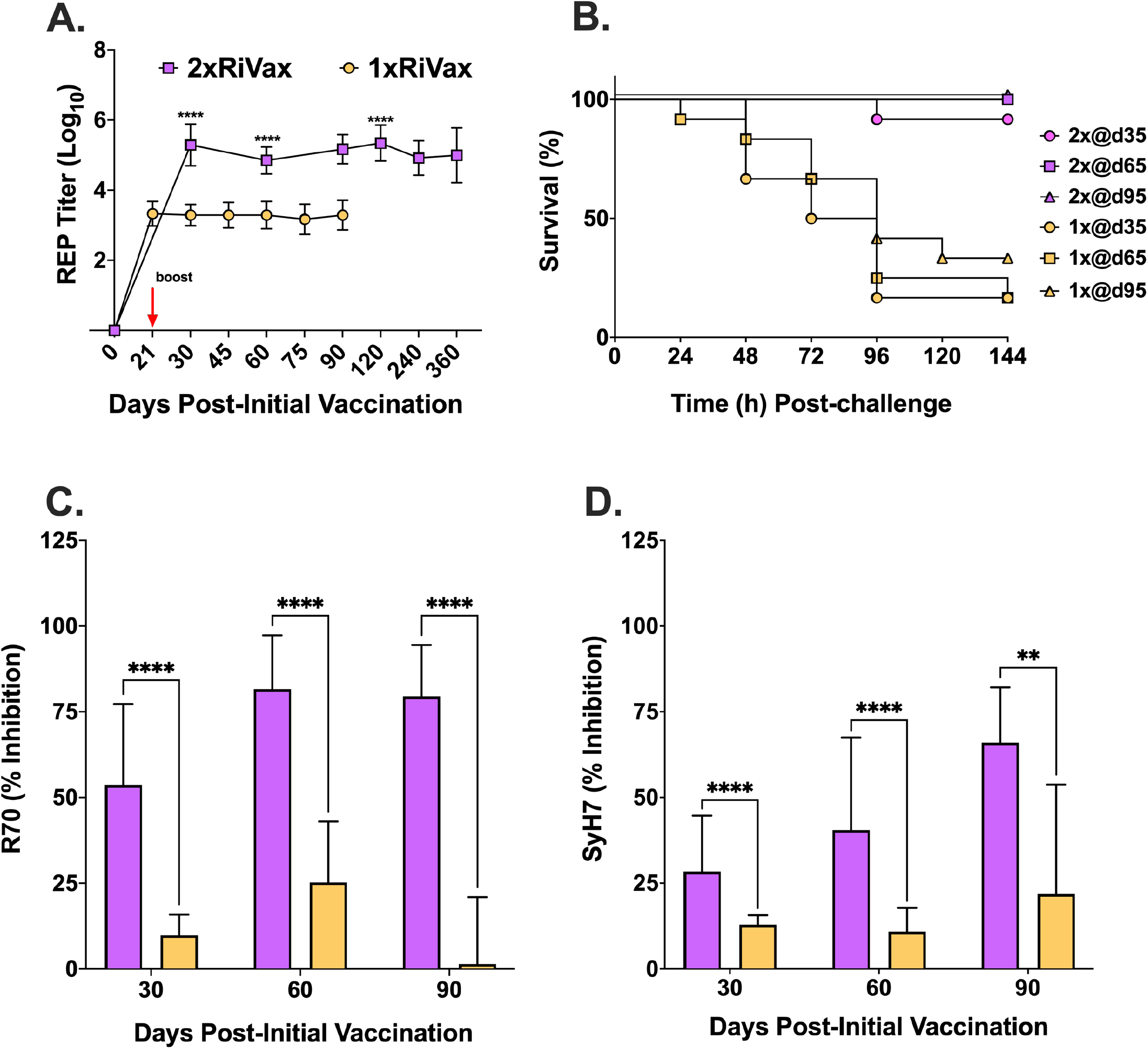
Comparison of single and two dose RiVax vaccinations on immunity to ricin toxin. Graphical representations of results presented in Table 2. (**A**) RT-specific reciprocal endpoint (REP) titers of serum IgG from mice vaccinated with 3 μg RiVax on day 0 (1 × RiVax; gold symbols) or on days 0 and 21 (2 × RiVax; purple symbols). (**B**) Kaplan-Meier plots of groups of RiVax-vaccinated mice challenged on study days 35, 65, and 95. (**C-D**) EPICC inhibition values from sera collected on study days 30, 60 or 90 from mice vaccinated with 3 μg RiVax once (gold) or twice (purple). A two-way ANOVA followed by a Šidák’s multiple comparisons test was used to determine significant differences in REP titers between groups, while mixed effects models followed by Šidák’s tests were used to examine differences in EPICC inhibition. The single mouse that died due to RT challenge on day 35 was excluded from this analysis. * P ≤ 0.05, ** P < 0.01, *** P < 0.001, **** P < 0.0001.

### Mucosal immunity to RT elicited by RiVax vaccination

RT is particularly toxic when inhaled (10, 22). While numerous reports have indicated that parenteral (systemic) RiVax vaccination is sufficient to confer immunity against a pulmonary RT challenge, those studies have been done with three doses, not two (19, 22). Therefore, cohorts of mice (n=12) were vaccinated SC with 0.3, 1, or 3 μg of RiVax on days 0 and 21. Mice were then challenged intranasally with a 10 × LD50 dose of RT on days 35, 125, and 162. Serum samples were collected from the animals five days prior to all the challenge days (plus an additional collection on day 90) and then evaluated by ELISA and EPICC (**Table 3**).

**Table 3.** Response elicited by RiVax prime-boost vaccination regimen followed by IN challenge.

|  |  |  |  |  |  |  |  |
| --- | --- | --- | --- | --- | --- | --- | --- |
| **Group** | **n=** | **Dose (µg)** | **RT (day)<sup>a</sup>** | **GMT<sup>b</sup>** | **% Inhibition (+/- SD)<sup>c</sup>** |  | **Survival<sup>d</sup>** |
|  |  |  |  |  | **R70** | **SyH7** |  |
| 1 | 12 | 0.3 | 35 | 3.130 | 15.09 (16.80) | 6.25(8.98) | 0/12 (0%) |
| 2 | 12 | 1 | 35 | 4.230 | 38.78 (25.98) | 20.09 (17.0) | 4/12 (33%) |
| 3 | 12 | 3 | 35 | 4.734 | 56.28 (25.05) | 28.4 (21.76) | 7/12 (58%) |
| 4 | 12 | 0.3 | 125 | 3.150 | 39.25 (31.49) | 24.43 (20.83) | 0/12 (0%) |
| 5 | 12 | 1 | 125 | 4.144 | 86.40 (15.77) | 67.12 (17.78) | 9/12 (75%) |
| 6 | 12 | 3 | 125 | 4.363 | 82.85 (18.06) | 60.75 (20.58) | 10/12 (83%) |
| 7 | 12 | 0.3 | 162 | 3.766 | 35.34 (39.44) | 33.62 (26.6) | 1/12 (8.3%) |
| 8 | 12 | 1 | 162 | 4.124 | 60.43 (33.34) | 64.23 (19.24) | 8/12 (67%) |
| 9 | 12 | 3 | 162 | 4.403 | 70.57 (24.26) | 62.6 (21.73) | 9/12 (75%) |
| 10 | 6 | N/A | 35 | 0 | 6.23 (3.34) | 8.62 (2.76) | 0/6 (0%) |
| 11 | 6 | N/A | 125 | 0 | 3.66(3.33) | 13.86 (3.27) | 0/6 (0%) |
| 12 | 6 | N/A | 162 | 0 | 6.04 (11.47) | 5.07 (4.11) | 0/6 (0%) |

<sup>a</sup>, RT challenge day ; <sup>b</sup>, geometric mean titers (log10); <sup>c</sup>, EPICC inhibition values, as described in the Materials and Methods; <sup>d</sup>, survivors/total mice per group over course of 7 days post RT challenge

Overall, mice that were vaccinated with 0.3, 1, or 3 μg of RiVax on days 0 and 21 fared poorly when challenged with RT intranasally on day 35. Indeed, only a single mouse that received the low dose of RiVax (0.3 μg) survived RT challenge (**Table 3**). In mice that received 1 μg of RiVax, survival ranged from 33% on day 35 to 67-75% on days 125 and 162. Groups of mice that received 3 μg of RiVax had 58% (7/12) survival on day 35, 83% (10/12) on day 125, and 75% (9/12) on day 162 (**Table 3**). The kinetics of RT-specific antibody responses were similar to those observed previously in that there were both dose- and time-dependent components to the onset of RT-specific serum IgG levels (**Table 3**). The 3 μg group displayed the highest titers and were significantly different from the 0.3 μg group at all time points, except day 157 (**Figure 4A**). R70 and SyH7 EPICC serum inhibition values were highest in the 3 μg RiVax group of mice (**Figure 4B, C**). Comparing groups of vaccinated mice by the same regimen but challenged by the IP versus IN routes suggests a higher serum IgG threshold associated with protection against 10 × LD_50_ RT mucosal challenge.

**Figure 4:**
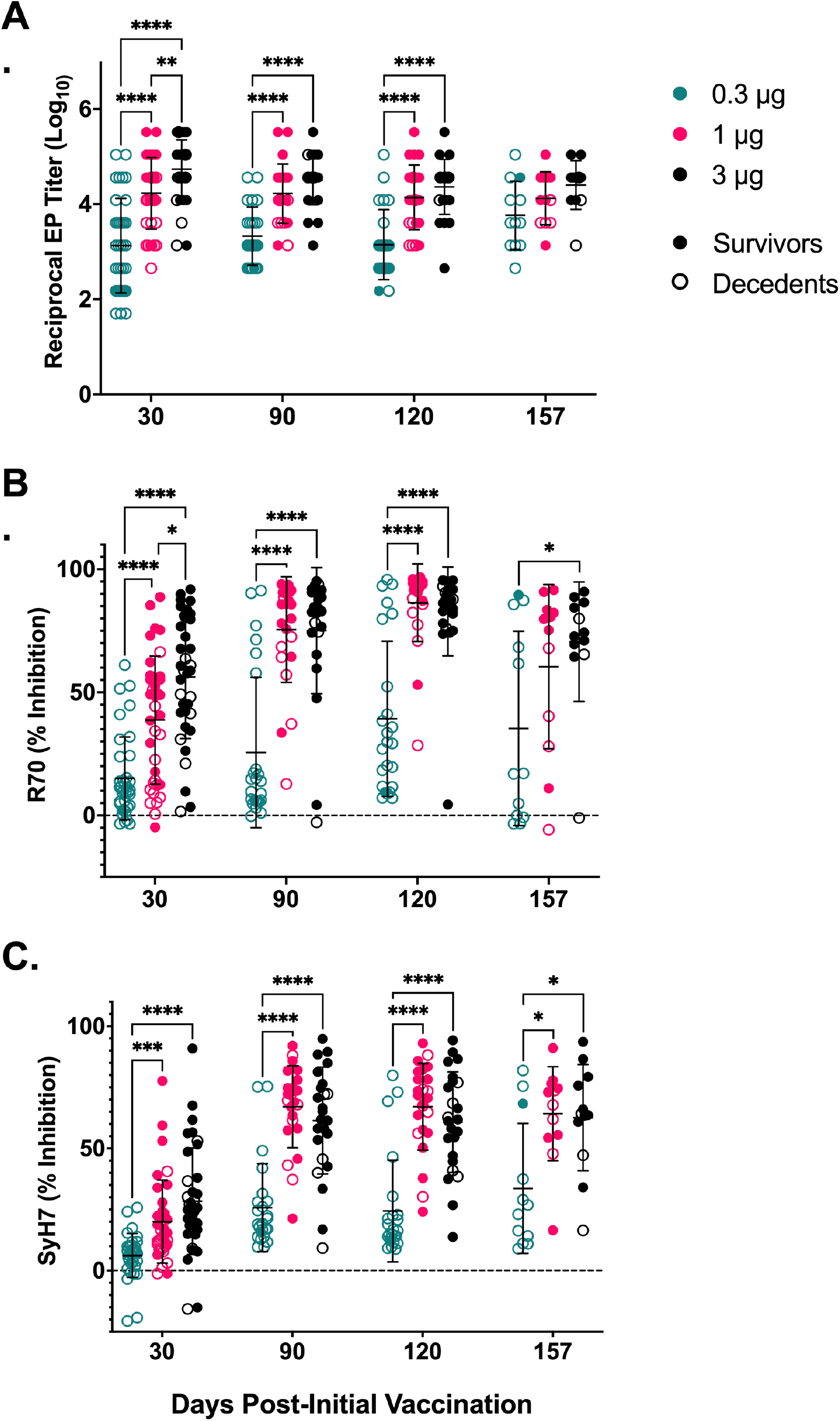
Serum antibody titers and EPICC inhibition values of R70 and SyH7 increased significantly with RiVax dose. Mice were vaccinated with 0.3, 1, or 3 μg of RiVax on days 0 and 21. EPT and EPICC inhibition values were determined for serum collected on days 30, 90, 120, and 157. Reciprocal endpoint titers were significantly higher in the 3 μg group than the 0.3 μg group at the 30, 90, and 120 timepoints, and were also significantly higher than the 1 μg group on day 30 (A). Similarly, R70 inhibition was significantly higher in the 3 μg than the 0.3 μg group at all timepoints, and higher than 1 μg group on day 30. Inhibition in the 1 μg group was significantly higher than the 0.3 μg group on days 30, 90, and 120 (B). Levels of SyH7 inhibition were also significantly higher in the 3 and 1 μg groups than the 0.3 μg group on all days (C). Mixed effects analyses followed by Šidák’s multiple comparisons tests were used to determine significant differences between groups. * P ≤ 0.05, ** P < 0.01, *** P < 0.001, **** P < 0.0001.

## DISCUSSION

The development of medical countermeasures against Select Agents and Toxins remains a research priority in the United States, even as the country is struggling to contain a resurgence of COVID-19. In this report we advanced pre-clinical studies of RiVax, a candidate RT subunit vaccine intended for use in military and high-risk civilian populations (25). We found that a parenteral prime-boost RiVax vaccination regimen in mice was sufficient to stimulate protective immunity to RT for at least 12 months. RT-specific serum IgG levels peaked between days 30 and 60, then persisted for the duration of the year-long study. Epitope-specific antibody responses, which are known to correlate with protective immunity to RT in mice, were equally durable (20). However, mice that received only a single dose of RiVax mounted significantly lower RT-specific serum IgG levels compared to their two-dose counterparts and fared poorly upon RT challenge, likely due to insufficient maturation of the toxin-neutralizing antibody response. In summary, we conclude that a prime-boost regimen of RiVax adjuvanted with Alhydrogel is both sufficient and necessary to elicit sustained immunity to parenteral RT challenge.

While the primary endpoint for assessing RiVax efficacy in this study was survival, secondary measures of toxin-induced morbidity, namely weight loss and hypoglycemia, provided insight into relative degrees of vaccine-induced immunity not evident from simple Kaplan-Meier plots (12, 21, 22, 26). For example, mice vaccinated with 3 μg of RiVax on days 0 and 21, then challenged on day 35, experienced significantly greater weight loss and hypoglycemia in the days immediately following toxin challenge than did an identically vaccinated group of mice challenged on day 365. This observation is consistent with a maturation of the RT antibody response over a period of >60 days, as has been noted previously in mice (27) and even non-human primates (19). In Rhesus macaques, for example, toxin-neutralizing antibody titers elicited following RiVax vaccination peaked several months after RT-specific IgG levels were maximal (19).

Another metric used in this study to interrogate the RT-specific antibody response following RiVax vaccination was EPICC, a competition ELISA with two toxin-neutralizing MAbs, R70 and SyH7, that target spatially distinct epitopes on RiVax (24). In a recent study, we demonstrated in a large cohort of RiVax-vaccinated mice that EPICC values combined with RT-specific endpoint titers in a multivariate model resulted in greater performance at predicting survival following lethal dose RT challenge than any one of the independent variables alone (20). EPICC inhibition values also serve as correlates of RiVax-induced protection in NHPs (C. Roy, D. Ehrbar, G. Van Slyke, E. Vitetta, O. Donini, N. Mantis, *manuscript in preparation*). While survival predictions were not necessarily applicable to the current study given the overall high survival rate, the kinetics of R70 and SyH7 EPICC inhibition values can be considered as readouts of RT-specific antibody maturation. For example, R70 EPICC inhibition values following two dose RiVax vaccination peaked on day 60, even though RT-specific endpoint titers peaked on day 30. In this respect, R70 EPICC inhibition values corresponded to the timepoint where full protection against RT challenge was achieved. SyH7 EPICC inhibition values peaked even later (day 120), possibly reflecting the subdominant nature of SyH7’s epitope, which is located on the backside of RTA relative to the active site (24). While we have not yet defined an absolute EPICC inhibition “threshold” associated with protective immunity to RT, the cumulative differences in R70 and SyH7 EPICC inhibition values between groups of vaccinated mice that survived RT challenge and those that did not are often stark, as exemplified in Figure 3.

Results from the RiVax studies involving parenteral vaccination followed by intranasal RT challenge reinforces the notion that the threshold of protection against RT respiratory exposure is greater than systemic exposure. In other words, higher levels of RT-specific serum IgG titers are required to protect mice against an intranasal (or aerosol) RT challenge, as compared to a parenteral (intraperitoneal) challenge. From passive MAb immunization studies in mice, we estimated that the threshold for immunity against intranasal RT challenge is about ∼8-10-fold greater than what was required to protect mice against the same amount of toxin given by intraperitoneal injection (28). We speculate that the higher serum IgG levels are due to issues related to lung bioavailability, as noted by others (29). Specifically, high serum IgG levels may be necessary to ensure sufficient transudation and/or transport of antibodies into the airways where they can intercept and neutralize RT before the toxin is internalized by alveolar macrophages and lung epithelial cells (28). The issue of serum IgG delivery into the lung has come to the forefront recently when considering SARS-CoV-2 vaccine efficacy and MAb therapy aimed at dampening viral replication and inflammation (30, 31). Of note, previous studies with aerosol exposure in Rhesus macaques have demonstrated that it is entirely feasible to obtain these higher levels of protection (19).

An investigation into the benefits of combining RiVax with an additional adjuvant may be warranted in light of emerging work on SARS-CoV-2 RBD subunit vaccines. Of particular note, there has been the success of aluminum salts combined with CpG in stimulating SARS CoV-2 neutralizing antibody titers in mice (32, 33). The aluminum salts plus CpG combination also stimulated unique cytokine and chemokine profiles in human PBMCs (33). The RiVax antigen is compatible with other adjuvants such LT-II (34, 35) and alpha-galactosyl ceramide (36), but, to our knowledge, RiVax adsorbed to aluminum hydroxide has not itself been combined with other adjuvants. Moreover, whether such adjuvants would be compatible with the RiVax lyophilization protocol would also need to be investigated (17), although success with a similar lyophilized formulation and a nano-emulsion adjuvant (CoVaccine HT™) has been demonstrated (37).

## MATERIALS AND METHODS

### Ricin toxin and RiVax vaccine formulations

Ricin (*Ricinus communis* agglutinin II) and RTA were purchased from Vector Laboratories (Burlingame, CA). Ricin was purchased without azide and sterility of the preparation was confirmed prior to use in animal studies. A single lot of lyophilized RiVax^®^ formulated as 10 mM histidine, 8% (w/v) trehalose, 200 μg/ml RiVax^®^, 0.85 mg/ml aluminum, pH 6.5) was used for all experiments (19). RiVax was reconstituted with sterile water for injection (1 mL) immediately prior to use.

### Mouse vaccination studies

Mouse studies were conducted under strict compliance with the Wadsworth Center’s Institutional Animal Care and Use Committee (IACUC). Female BALB/c mice aged 6-8 weeks were obtained from Taconic Biosciences (East Greenbush, NY) and housed under conventional, specific-pathogen-free conditions. Mice (n = 12 per experimental group; n = 6-8 for control groups) were vaccinated via the subcutaneous (SC) route on days 0 and 21. The vaccines were administered to mice on days 0 and 21 in 50 μl final volumes. Blood was collected from mice via the submandibular vein on days indicated in the figures.

### RT challenge

Animal experiments were conducted in compliance with the Wadsworth Center’s Institutional Animal Care and Use Committee (IACUC). The Wadsworth Center is an AAALAC accredited institution. Mice were challenged by intraperitoneal (IP) injection or intranasal instillation (under isoflurane anesthesia) with the equivalent of 10 × LD_50_ (100 μg/kg) RT. Survival was monitored over a 7-day period. Blood glucose levels were measured for four days post challenge, and body weight daily until day 7. As per protocol, mice were euthanized when they became overtly moribund, experienced weight loss ≥ 20% of pre-challenge weight, and/or became hypoglycemic (<25 mg/dl) following IP challenge or (<49 mg/dl) for IN challenge.

### ELISA

Nunc Maxisorb F96 microtiter plates (ThermoFisher Scientific, Pittsburgh, PA) were coated overnight at 4 °C with ricin (1 μg/well in PBS), washed with PBS with 0.05% (v/v) Tween-20 (PBS-T), and then blocked for 2 h with PBS-Tween (PBST) containing 2% (v/v) goat serum (Gibco, MD, USA). Three-fold serial dilutions of serum (starting at 1:50) were then applied to plate for 1 h at ambient temperature, washed and detected with horseradish peroxidase (HRP)-conjugated goat anti-mouse IgG (SouthernBiotech, Birmingham, AL). The ELISA plates were developed using SureBlue 3,3′,5,5′-tetramethylbenzidine (TMB; Kirkegaard & Perry Labs, Gaithersburg, MD) and analyzed using a SpectroMax 250 spectrophotometer equipped with Softmax Pro 5.4.5 software (Molecular Devices, Sunnyvale, CA). The endpoint titer was defined as the minimal dilution whose absorbance (450 nm) was > 3 times background, with background being defined as the average absorbance produced by wells with buffer alone. Seroconversion was defined as the endpoint titer of ≥ 1:50. Geometric mean titers (GMTs) were calculated from the endpoint titers. Mice that had not seroconverted, as determined by ELISA, were assigned a GMT of 1 for the purposes of statistical analysis. Vero cell cytotoxicity assays were performed as previously described (38, 39).

### EPICC analysis

Epitope profiling immunocapture competition (EPICC) assays were conducted as described (20). Immulon 4HBX 96 well plates (ThermoFisher Scientific, Pittsburgh, PA) were coated overnight with a capture anti-RTA mAb (1 μg/ml). The following day the plates were blocked with 2% goat serum, washed, and then overlaid with a mixture of biotin-tagged ricin and polyclonal antibody (pAb) mouse serum (1:25). The amount of biotin-ricin used in the competition ELISA was equivalent to the EC_90_ for each capture mAb (range: 100-150 ng/ml). After 1 h incubation, plates were washed and developed with streptavidin-HRP (1:1000; ThermoFisher Scientific, Pittsburgh, PA) and TMB, as described above for ELISAs. The percent (%) inhibition of ricin binding to the capture mAb in the presence of pAb serum was calculated from the optical density (OD) values as follows: 1 – value OD_450_ (biotin-ricin + competitor mAb) ÷ value OD_450_ (biotin-ricin without competitor mAb) × 100.

### Statistical analysis

Statistical analyses was performed with GraphPad Prism v. 8.01. Reciprocal endpoint titers were log-transformed prior to statistical analysis. Two-way ANOVAs or mixed effects models followed by Šidák’s multiple comparisons tests were performed to determine significant differences between reciprocal endpoint titers and EPICC inhibition levels, as described in the Figure legends. Survival data were tested using the log rank Mantel-Cox test. In all cases, the significance threshold was set at P < 0.05.

## Acknowledgements

We thank Greta Van Slyke (Wadsworth Center) for valuable feedback and guidance. We gratefully acknowledge the Wadsworth Center’s Veterinary Sciences staff for maintenance of RiVax-vaccinated mice during COVID-19 imposed work hours. This study was supported by grant AI125190 (to NJM) and Contract No. HHSN272201400039C (to OD; Soligenix, Inc.) from the National Institutes of Allergy and Infectious Diseases, National Institutes of Health (NIH). The content is solely the responsibility of the authors and does not necessarily represent the official views of the NIH. The funders had no role in study design, data collection and analysis, decision to publish, or preparation of the manuscript.

## Author Contributions

HN and JD conducted the experiments; HN, JD, DE, OD and NJM analyzed the results; HN, JD, and NJM wrote the manuscript.

